# Excessive chondrogenesis in the blastema initiates during hypomorphic limb regeneration in *Xenopus* froglet, but not during patterned limb regeneration in *Xenopus* tadpoles and newts

**DOI:** 10.1101/2025.03.29.646070

**Authors:** Suzu Kobari, Hibiki Yokoyama, Kagayaki Kato, Rina Sato, Nana Kitagawa, Joe Sakamoto, Yasuhiro Kamei, Hitoshi Yokoyama

**Affiliations:** Department of Biochemistry and Molecular Biology Faculty of Agriculture and Life Science, Hirosaki University, 3 Bunkyo-cho, Hirosaki, Aomori 036-8561, Japan; Optics and Imaging Facility, Trans-Scale Biology Center, National Institute for Basic Biology, Myodaiji, Okazaki, Aichi 444-8585, Japan; Biophotonics Research Group, Exploratory Research Center on Life and Living Systems (ExCELLS), National Institute for Physiological Sciences, Higashiyama Myodaiji, Okazaki, Aichi 444-8787, Japan; Department of Basic Biology in the School of Life Science of the Graduate University for Advanced Studies (SOKENDAI), Okazaki, Aichi 444-8585, Japan

**Author notes:** Author for correspondence: Hitoshi Yokoyama, Department of Biochemistry and Molecular Biology, Faculty of Agriculture and Life Science, Hirosaki University, 3 Bunkyo-cho, Hirosaki, Aomori 036-8561, Japan.

**Keywords:** limb regeneration, cartilage, chondrogenesis, Sox9, *Xenopus*, newt

## Abstract

*Xenopus laevis* tadpoles and newts regenerate a limb almost completely after amputation, with recapitulation of the patten formation of the limb. Metamorphosed *Xenopus* froglets form a cone-shaped regenerating blastema, similar to tadpoles and newts, but ultimately regenerate only a hypomorphic cartilaginous spike. Previous study suggested that excessive chondrogenesis, distinct from *Xenopus* tadpoles, may occur in the regenerating limb of a froglet and may prevent pattern formation during regeneration. However, it remains unclear whether excessive chondrogenesis actually occurs in froglet blastemas. If it does, when does it initiate and how does it progress in the blastemas? To answer these questions, we examined the extent of chondrogenesis in regenerating blastemas which have the common morphological shapes observed in newts (*Pleurodeles waltl*), *Xenopus laevis* tadpoles, and froglets. To evaluate excessive chondrogenesis, we developed a simplified procedure using immunofluorescence for cartilage markers (Sox9 or Col2a1) and quantitative image analysis. Our analysis revealed that signs of excessive chondrogenesis were detected not in newts and tadpoles but in froglets blastemas. During limb regeneration in froglets, the first sign of excessive chondrogenesis was detected in the cone-shaped blastema at the medium bud (MB) stage, and excessive chondrogenesis progressed to a more severe state as the blastema grew. These results indicate that excessive chondrogenesis initiates specifically in froglet blastemas at the MB stage at the latest and progresses in a definite spatio-temporal manner. Further elucidation of the mechanisms underlying froglet-specific excessive chondrogenesis in the blastema may lead to the recovery of patterned limb regeneration in froglets with adequate inhibition of chondrogenesis.

## INTRODUCTION

Newts are often called the champions of regeneration among adult vertebrates since they regenerate amputated limbs, transected ventricles (hearts), removed lenses (eyes), and so on throughout their life cycles (Straube and Tanaka, 2006; Joven et al., 2019). In the case of limb regeneration, a newt regenerates a well-patterned limb along the three primary limb axes—anteroposterior (AP), dorsoventral (DV), and proximodistal (PD) axes (Otsuki et al., 2022). Anuran amphibians also regenerate amputated limb almost completely when their limb primordia (limb buds) are amputated at early tadpole stages but their regenerative capacity for limb declines as metamorphosis proceeds (Stocum, 1995). For example, in *Xenopus laevis*, the most widely used anuran model, early-stage tadpoles prior to metamorphosis regenerate a well-patterned limb after amputation, but the froglet, a young adult soon after metamorphosis, regenerates only a hypomorphic limb (Fig. 1A; Dent, 1962; Robinson and Allenby, 1974). The hypomorphic limb regenerate, called a “spike”, is an unsegmented, rod-shaped structure composed exclusively of cartilage without muscle (Fig. 1B; Dent, 1962; Endo et al., 2000; Robinson and Allenby, 1974; Satoh et al., 2005; reviewed by Suzuki et al., 2006). While it regenerates a hypomorphic limb, an amputated limb of a froglet undergoes common processes with that of a newt (urodele amphibian)—the wound epidermis covers the amputation plane quicky, and then blastema formation occurs in a nerve-dependent manner (Suzuki et al., 2006; Stoick-Cooper et al., 2007; Joven et al., 2019). Limb regeneration in a froglet seems to fail in pattering along the AP, DV and PD axes and in re-differentiation such as muscle differentiation, after blastema formation (Endo et al., 2000; Matsuda et al., 2001; Satoh et al., 2005; Yakushiji et al., 2007; Ohgo et al., 2010). However, the original cause(s) for the failure in patterning and re-differentiation during limb regeneration in a froglet remains mostly unclear.

**Fig. 1.**
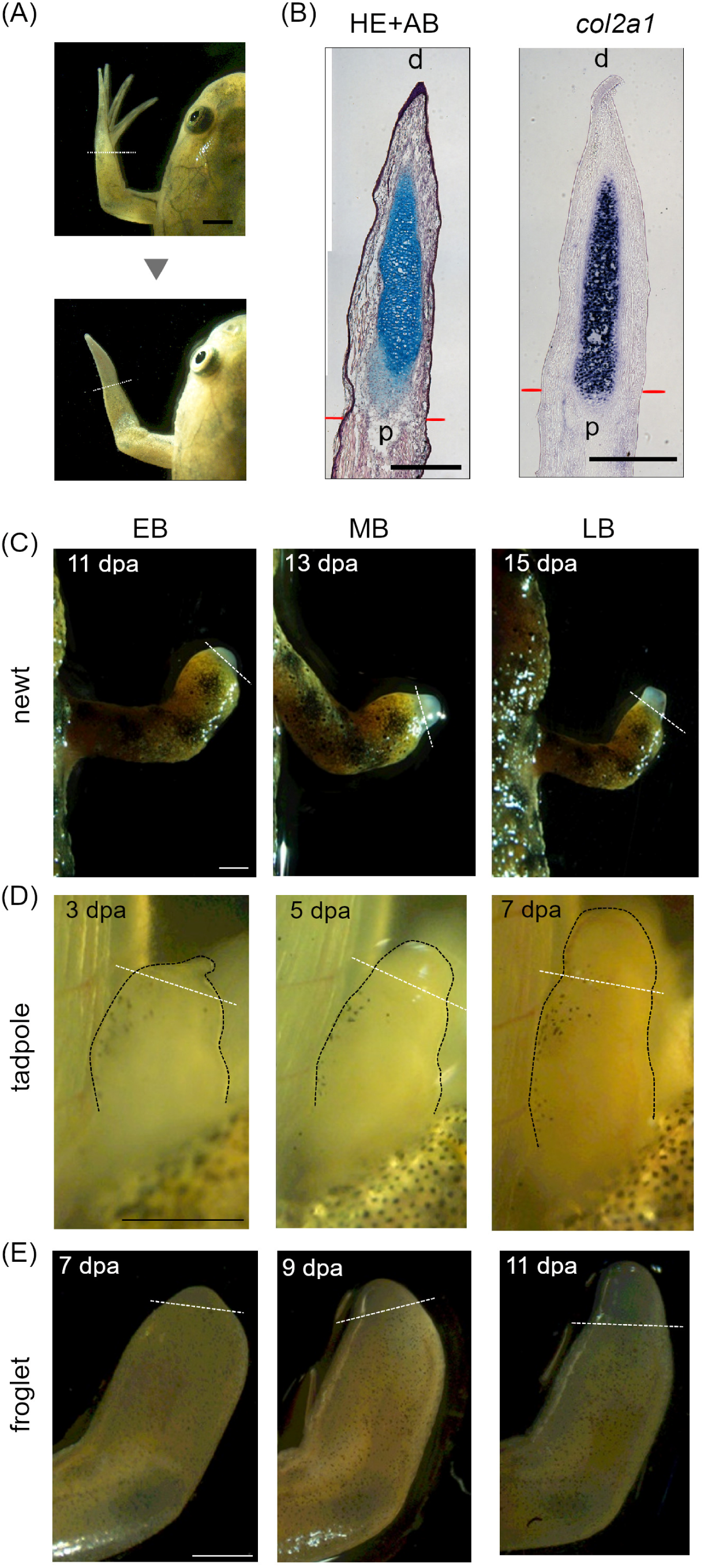
Cartilaginous spike regeneration in a *Xenopus* froglet and stages of amphibian limb regeneration. (A) Limb amputation in a froglet resulted in hypomorphic limb regeneration at 28 dpa. (B) Longitudinal adjacent sections of a spike regenerate at 28 dpa were underwent hematoxylin, eosin and alcian blue (HE+AB) staining or *in situ* hybridization with a *col2a1* antisense probe. d, distal; p, proximal. Newt (C), tadpole (D) and froglet (E) blastemas at the EB, MB and LB stages. Note that blastemas and stumps are outlined dotted black lines in (D). A dotted white line or a pair of red lines, the amputation level. Bars = 2 mm for (A), 400 μm for (B) and 1 mm for (C, D, E).

As the putative cause of the failure in patterning and re-differentiation, Ohgo et al. (2010) proposed that premature and excessive differentiation of blastema cells into chondrocytes (cartilage cells) may prevent pattern formation along the axes and muscle formation. In chondrogenesis (differentiation into cartilage cells), Sox9 is an essential transcription factor for this process and its expression is an early marker for differentiating chondrocytes (Egawa et al., 2014; Kozhemyakina et al., 2015). Sox9, along with two additional Sox family members (Sox6 and Sox6), cooperatively promotes proliferation and differentiation of chondrocytes by activating the expression of genes encoding extracellular matrix (ECM) proteins. These ECMs, encoded by the direct target genes of Sox9, include collagen type II (Col2a1) and Aggrecan (Acan), which play critical roles in chondrocyte differentiation (Egawa et al., 2014; Kozhemyakina et al., 2015). In limb regeneration in *Xenopus*, gene expression of *sox9* was detected by *in situ* hybridization in the regenerating limb blastema. The *sox9* expression domain appeared to be broader in the froglet blastema than in the tadpole blastema (Ohgo et al., 2010). Furthermore, quantitative RT-PCR revealed that the amount of *sox9* transcript was significantly more abundant in the froglet blastema at 14 days post-amputation (dpa) than in the tadpole blastema at 7dpa (Ohgo et al., 2010). Based on these results, Ohgo et al. (2010) proposed that premature and excessive differentiation of blastema cells into chondrocytes may be responsible for failure of pattern formation and muscle regeneration during limb regeneration in froglets.

However, to draw a conclusion regarding premature differentiation into chondrocytes, several points remain to be clarified. Firstly, the tadpole and froglet blastema need to be compared at the same (or comparable) stage of limb regeneration. While qRT-PCR analysis (Ohgo et al., 2010) detected significantly higher expression of *sox9* in the froglet blastema at 14 dpa than in the tadpole blastema at 7 dpa, the blastema stages, defined by their morphological shape, appear to differ between the tadpole and froglet. Secondly, qPCR quantifies gene expression abundance but does not prove spatial information (spatial pattern) about gene expression. Thus, we cannot conclude that the blastema is widely populated by chondrocytes based solely on qPCR data. Thirdly, it is important to compare not only between tadpole and froglet within the species *X. laevis*, but also between froglets (*X. laevis*) and young adult newts soon after metamorphosis (referred to simply as “newts” hereafter) to test the possibility of excessive chondrogenesis in the froglet blastema. Fortunately, *Pleurodeles waltl* has become increasingly available in the amphibian research community as a new model urodele since the 2010s (Hayashi et al., 2013; Matsunami et al., 2019). To address these issues, this study aims to compare the blastemas of tadpoles, froglets, and newts at the same regeneration stage, analyzing the spatial distribution of Sox9-expressing cells within the blastema, and investigating whether excessive chondrogenesis occurs in the froglet blastema.

To compare the blastemas of tadpoles, froglets, and newts at comparable stages of limb regeneration, we used stages of limb regeneration in *Notophthalmus viridescens* (Eastern spotted newts) as described by Iten and Bryant (1973). Blastemas at early bud (EB), medium bud (MB), and late bud (LB) stages were included in the following comparative analysis. After amputation, the wound epidermis quickly covers the amputation site, and blastema cells accumulate under the wound epidermis in all tadpoles, froglets, and newts. We specifically focused on the EB and MB stages, where the blastema appears dome-shaped and cone-shaped, respectively. While the blastema can be easily identified at the amputation site, asymmetrical polarity along the AP or DV axis is not apparent at these stages. The morphological shape of blastema is similar among tadpoles, froglets, and newts. Thus, if excessive chondrogenesis is detected specifically in froglets at these stages by analyzing the spatial distribution of Sox9-expressing cells, this could be an important finding for predicting the fate of limb regeneration in froglets versus tadpole and newts, particularly in the context of hypomorphic and patterned limb regeneration.

To evaluate excessive chondrogenesis with spatial information, we performed immunofluorescence against Sox9 protein on the blastema sections and analyzed the distribution of differentiating chondrocytes. Through intraspecific (tadpole versus froglet) and interspecific (newt versus froglet) comparisons, we assessed chondrogenesis in the blastemas at the same stages of regeneration. We defined a rectangular-shaped area of analysis, extending from the proximal base to the apical (distalmost) tip of the blastema. In the distal-to-proximal direction, we examined whether the Sox9 and Col2a1 (a typical downstream factor of Sox9 in chondrogenesis) signals increased significantly and sharply at a specific point (change point) along the PD axis using an R package. A higher incidence of change point detection and a more distal location of change point indicate more severe excessive chondrogenesis in the blastema. In this analysis of the blastemas from tadpoles, froglets and newts, the change point was specifically detected in the froglet blastema. The change point of the Sox9 signal was first detected at the MB stage, while the change point of the Col2a1 signal was firstly detected at the subsequent stage (LB). These results suggest that signs of premature and excessive chondrogenesis initiate not in the tadpole or newt blastema but in the froglet blastema, at the latest by the MB stage.

## MATERIALS AND METHODS

### Animal husbandry and limb amputation

Wild-type *Xenopus laevis* adult frogs were purchased from domestic animal venders, Watanabe Zoushoku (http://www5d.biglobe.ne.jp/~zoushoku/top.htm) and Hamamatsu Seibutsu Kyozai (http://www.h-seibutsu.co.jp/index.html). Fertilized eggs were obtained by natural mating. Tadpoles and froglets were reared at 22–23°C in dechlorinated tap water. The rearing containers were cleaned daily, and the tadpoles were fed powdered barley grass (Odani Kokufun, Japan or Yamamoto Kanpoh, Japan). At stage 58 (Nieuwkoop and Faber, 1994), the feeding was ceased until metamorphosis was completed. After metamorphosis, the froglets were fed dried tubifex every other day.

Wild-type adult newts (*Pleurodeles waltl*) were provided by Hiroshima University Amphibian Research Center through National BioResource Project (NBRP) of MEXT. Fertilized eggs were obtained by artificial insemination according to Hayashi et al. (2013). Larval and juvenile newts were reared at 22–26°C in dechlorinated tap water. They were fed hatched brine shrimp daily until 7 weeks after fertilization, after which they were switched to combined feed (Kyorin Cooperation, Hyodo, Japan).

For limb amputation for blastema formation, *Xenopus* tadpoles, froglets and juvenile newts were anesthetized with 0.05% ethyl-3-aminobenzoate (Tokyo Chemical Industry, 886-86-2) dissolved in Holtfreter’s solution. For froglets and juvenile newts, both of which had recently completed metamorphosis, their forelimbs were amputated through the distal zeugopodium using Noes scissors (straight) blade 10mm (Nisshin EM Co., Ltd., Japan), and the amputation surface was trimmed to make it flat. For *Xenopus* tadpoles, hindlimb buds at stage 53 were amputated at the presumptive ankle level (according to the fate map by Tschumi, 1957) using Noes scissors (straight) 5mm (Nisshin EM Co., Ltd., Japan). Formed blastemas were observed under a stereomicroscope (Nikon SMZ18) with a 0.5x objective lens and photographed by a digital camera (Olympus DP74). All animal care and experimentation procedures were conducted in accordance with the animal care and use committee guidelines of Hirosaki University (A15003, A15003-1).

### Immunofluorescence

The limb blastemas (for immunofluorescence) were excised and fixed for 2 h at 4°C in 4% paraformaldehyde/phosphate-buffered saline (PFA/PBS), decalcified overnight at 4°C with 8% ethylenediaminetetraacetic acid (EDTA)/PBS, and embedded in Optimal Cutting Temperature (OCT) compound (Sakura). Samples were then sectioned at 10 µm by a cryostat. The sections were mounted on PLATINUM PRO or MAS glass slides (Matsunami Glass).

For immunostaining, we used the protocol described by Suzuki et al. (2007). Briefly, frozen section samples were rinsed with wash buffer (PBS plus 0.02% Tween-20) three times for 10 min each. For type II collagen (Col2a1) staining, samples were incubated with 1 μg/ml Proteinase K/PBS at 37 °C for 15 min to induce antigen retrieval, rinsed with wash buffer three times for 1 min each, re-fixed with 4% PFA/PBS at room temperature for 20 min and rinsed again with wash buffer two times for 1 min each. After rinse with wash buffer, samples were incubated with primary antibodies diluted in blocking buffer (PBS plus 0.02% Tween-20, 0.1% Triton X-100 and 1% heat-inactivated goat serum) at 4 °C overnight. Primary antibodies were mouse anti-Msx1/2 monoclonal antibody (Developmental Studies Hybridoma Bank, 4G1-c, diluted 1:50), rabbit ant-Sox9 polyclonal antibody (Merck Millipore, AB5535, diluted 1:500) and mouse anti-Col2a1 monoclonal antibody (Developmental Studies Hybridoma Bank, II-II6B3, diluted 1:50). After the primary antibody incubation, sections were washed with wash buffer several times then incubated with secondary antibodies diluted in blocking buffer for 3 h at room temperature. Secondary antibodies were Alexa488 goat anti-mouse IgG1 (Thermofisher, A21121, diluted 1:1000), Alexa555 goat anti-rabbit IgG (Thermofisher, A21428, diluted 1:1000) and Alexa594 goat anti-mouse IgG(H+L) (Thermofisher, A11032, diluted 1:100). Sections were counterstained with DAPI (WAKO, 049-18801, diluted 1:20000) to detect nuclei, washed several times with wash buffer, and mounted in Fluoro-KEEPER Antifade Reagent, Non-Hardening Type (nacalai tesque, 12593-64).

### *in situ* hybridization and histological (hematoxylin, eosin and alcian blue; abbreviated as HE+AB) staining

Regenerating limbs (for *in situ* hybridization and histological staining) were fixed in MEMFA (0.1 M MOPS at pH 7.4, 2 mM EGTA, 1 mM MgSO_4_, 3.7% formaldehyde) overnight at room temperature, embedded in OCT compound (Sakura), and serially sectioned at 10 µm by a cryostat. For synthesis of probes for *in situ* hybridization, DIG-labeled RNA probes of *col2a1* (Matsubara et al., 2023a) were prepared according to the protocol of the manufacturer (Roche). *In situ* hybridization on frozen sections was performed according to Yoshida et al. (1996) with slight modifications. Sections were observed under an upright microscope (Olympus BX53) with an objective lens (Olympus UPlan PLN 10x) and were photographed by a digital camera (Leica DFC7000T).

For histological staining, sections were washed with PBS, deionized water and 70% ethanol for 5 min each. The sections were then stained with 0.1% Alcian blue (8GX, Merck Millipore, 1.05234) in 70% ethanol for 1 h at room temperature. After confirmation of sufficient staining under inverted microscope, the sections were washed several times with 70% ethanol until the ethanol became colorless. The sections were gradually rehydrated with 50% ethanol, 20% ethanol, and deionized water and then washed with running tap water. The sections were stained with Mayer Hematoxylin Solution (Wako 131-09655) for 40 min at room temperature and washed with running tap water for 1h. After confirmation of sufficient nuclear staining under inverted microscope, the sections were stained with 0.25% acidic eosin (Merck Millipore, 1.15935) in 60% ethanol for 30 seconds and washed with running tap water. To obtain optimal staining color for the eosin, the sections were repeatedly washed with 70% ethanol and then running tap water several times. The sections were dehydrated with 70%, 95% and 99.5% ethanol. Then the sections were cleared with xylene three times for 5 min each. Finally, the sections were sealed with EUKITT mounting medium (O. Kindler and ORSAtec).

### Imaging of section samples

For immunofluorescence, all blastemas were sectioned longitudinally across the anteroposterior (AP) axis. The immunostained section samples were observed under an FV3000 confocal laser scanning microscope (Olympus) equipped with an Olympus UplanSApo 10x or 20x dry objective lens. Fluorescence was detected using 405 nm, 488 nm or 561 nm excitation through a GaAsP photo multiplier tube. For subsequent “change point” analysis, two representative sections were selected, transected at the midpoint along the dorsoventral (DV) axis for each blastema sample. To select the representative sections, the two non-adjacent sections with the largest width along the AP axis from all available serial sections of a blastema. Optical-section images of the representative sections were then photographed for subsequent analysis.

### Change point analysis and image processing

Blastema images from the two representative sections of each blastema, as described above, captured by confocal microscopy, were analyzed using ImageJ (Fiji) (Fig. 2). To remove noise, all images were processed using a mean filter. The anterior and posterior edges between the regenerating blastema and the remaining proximal limb tissues in the image were defined based on both morphological shape and the immunostaining signals using an anti-Msx1/2 (a blastema marker) antibody (Fig. 2A, D). A straight line was then drawn connecting the two edges, serving as the boundary between the blastema area and the proximal limb tissues. The blastema area, excluding the epidermis, was cropped using the polygon area selection tools. Next, the length of the boundary line between the blastema and proximal tissues was measured in each section sample, and a rectangle-shaped ROI (region of interest) was defined by two vertical lines extending from the apical tip of the blastema to the boundary line (Fig. 2B, C, bottom). The vertical lines were positioned to cover the longest length, with the interval of approximately 20% of the full-length boundary line. If 20% of the full-length boundary exceeds ImageJ’s maximum width (300 pixels), the entire image was geometrically reduced until the length was below 300 pixels.

**Fig. 2.**
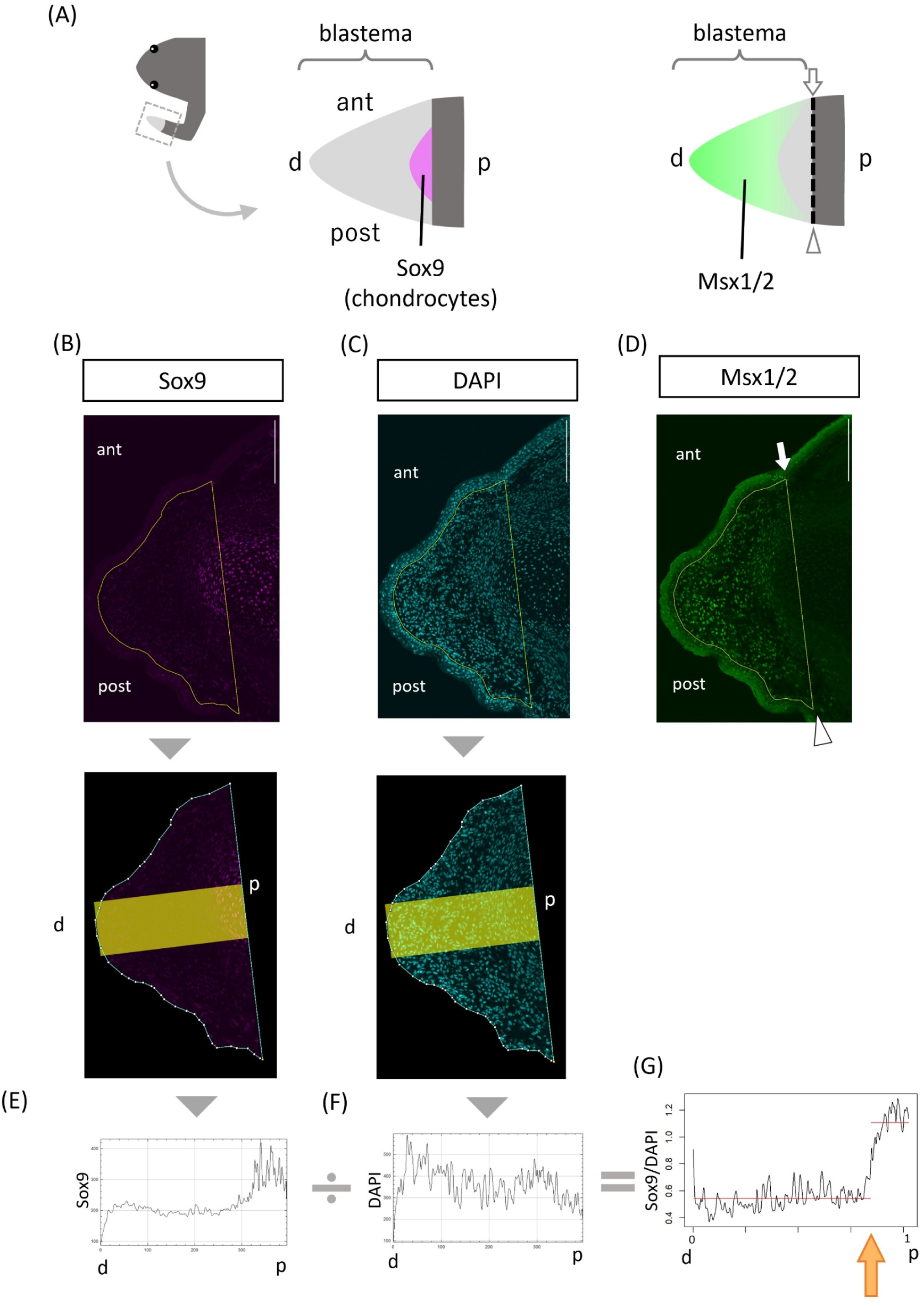
Change point analysis of immunofluorescence images for detection of excessive chondrogenesis in a blastema. (A) Schematic of change point analysis. The anterior and posterior edges of the blastema were defined by a blastema marker, Msx1/2 and the outline shape of the longitudinal section (left). The distribution of chondrocytes was visualized by an early chondrocyte marker, Sox9 (right). (B) Sox9, (C) DAPI, and (D) Msx1/2 images of a representative section of a froglet blastema at the MB stage. A yellow rectangle indicates the region of interest (ROI) for analysis. (E) Sox9 and (F) DAPI plot profiles from the ROI shown in (B) and (C), respectively. (G) Adjusted Sox9/DAPI plot profile acquired from the ROI shown in (B) and (C). The Sox9 profile was divided by the DAPI profile. An orange arrow indicates a detected change point. d, distal; p, proximal; ant, anterior; post, posterior; white arrow, anterior edge of blastema; arrowhead, posterior edge of blastema. Note that the brightness of immunofluorescence images has been increased from their original versions to improve visibility.

From the rectangular ROI, the plot profile of immunostaining signal with anti-Sox9 (Sox9 plot profile) or anti-Col2a1 antibody (Col2a1 plot profile) was acquired. Simultaneously, the corresponding plot profile of DAPI staining (DAPI plot profile) was acquired from the same ROI. The Sox9 or Col2a1 profile was divided by the corresponding DAPI profile to adjust for the effect of cell density (Fig. 2E, F, G), as the DAPI profile reflects cell density. To examine whether the rectangular ROI contains a position where the cartilage marker (Sox9 or Col2a1) signal significantly increases from distal to proximal, we detected the change point (cpt.mean) using *changepoint* package (v2.2.4) in R (v4.3.1) (Killick and Eckley, 2014). Change point value was represented as the relative level along the proximodistal (PD) axis, ranging from zero (the distal-most apical tip of the blastema) to 1.0 (the boundary between the blastema and proximal tissues) (Fig. 2G). If the change point was not detected in either of the two representative sections of a blastema, we judged that the blastema did not have a change point (in which case the value would be 1.0). If the change point was detected in at least one of the two representative sections, we judged that the blastema had a change point. If both sections indicated the change point, we collected only the change point value at the more distal level for the blastema sample.

Immunofluorescence signals in the representative images sometimes showed low brightness and contrast, making it difficult to visually identify convincing signals with the naked eye. To enhance the brightness and contrast in the representative images of immunofluorescence shown in Fig. 3 and Figs. S2-S5, image processing was performed. However, to ensure appropriate image manipulation, we adhered to the standard image processing guidelines outlined in scientific journals (e.g., https://www.nature.com/nature-portfolio/editorial-policies/image-integrity). ImageJ was used for this processing: the brightness and contrast were adjusted to make the signals more visible. Additionally, a calibration bar showing the range of grey values was added to each image using ImageJ. It is important to note that all change point analyses were performed using the original, unprocessed immunofluorescence images, not the processed versions.

**Fig. 3.**
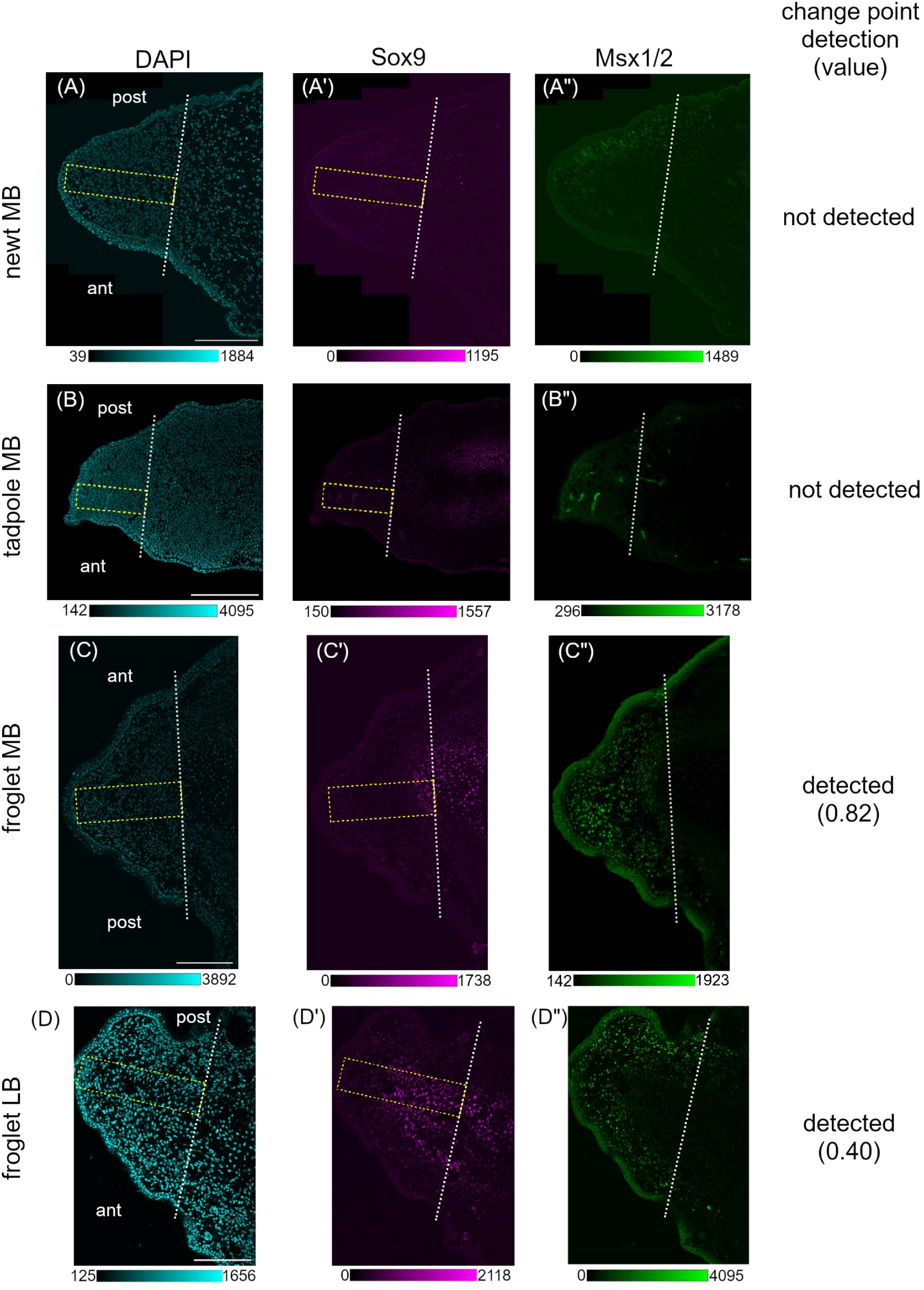
Representative immunofluorescence images of a blastema for change point analysis using anti-Sox9 antibody in a newt, tadpole and froglet. Fluorecence images of a longitudinal section of newt MB (A-A’’), tadpole MB (B-B’’), froglet MB (C-C’’, identical to Fig. 2C, B, D) and froglet LB blastemas (D-D’’) at the MB or LB stage. A yellow rectangle indicates the region of interest (ROI) for analysis. A dotted white line indicates the amputation level. A calibration bar, created using ImageJ, represents the range of grey values in each image. ant, anterior; post, posterior. Bars = 400 μm for (A) and 200 μm for (B, C, D).

## RESULTS AND DISCUSSION

### Limb regeneration in froglets results in a hypomorhic spike, but it undergoes common processes with limb regeneration in newts and tadpoles

A froglet’s limb regenerates not into a patterned limb but into a hypomorphic, spike-shaped limb after amputation (Fig. 1A; Dent, 1962; Robinson and Allenby, 1974). Once a regenerate reaches the late stage of regeneration and becomes rod-like in shape, it is composed exclusively of chondrocytes that are positive for typical cartilage markers (Fig. 1B; Satoh et al., 2005). While the final regenerate greatly differs from those of newts and tadpoles, limb regeneration in froglets undergoes common processes up until the formation of the cone-shaped blastema. For instance, all limbs of newts, tadpoles, and froglets initially form a dome-shaped early bud (EB) blastema, which then transitioned to a cone-shaped medium bud (MB) blastema at the amputation site, though the time taken to reach these blastema stages differs between species (Fig. 1C-E). After the MB stage, the blastemas of newts and tadpoles grow into late bud stage as defined by Iten and Bryant (1973), but those of froglets transform into a different shape. The froglet blastema elongates along the PD axis, similarly to newts and tadpoles, but does not flatten along the DV axis, while the newt and tadpole blastemas flatten at the LB stage. We expediently defined the elongated but non-flattened blastema as froglet “late bud (LB)” blastema (Fig. 1E, right). We specifically focused on the EB and MB stages for the following comparative analysis to identify the first signs of excessive chondrogenesis, as the fate of the froglet blastema appears to diverge from that of the newt and tadpole blastema at the LB stage.

### Comparative analysis to assess excessive chondrogenesis among newt, tadpole, and froglet using common criteria**—**change point analysis

To evaluate excessive chondrogenesis in regenerating limb blastemas, we performed immunofluorescence against Sox9 protein, the early marker for chondrogenesis as previously mentioned (Egawa et al., 2014; Kozhemyakina et al., 2015), on the longitudinal sections of blastema at the comparative stage. The same section was immunostained against Msx1/2 protein to visualize the blastema cells as described by Suzuki et al. (2007). Based on both the morphological shape and the Msx1/2 immunofluorescence signal, the anterior and posterior edges of the blastema area in each section were determined (Fig. 2A, D). A straight line was then drawn connecting the two edges, marking the boundary between the blastema area and more proximal tissues (see Materials and Methods for details.). A rectangle-shaped area was set from the apical tip to the boundary line around the midline along the AP axis (Fig. 2B, C, bottom).

For this rectangle area, the Sox9 immunofluorescence signal profile (Fig. 2E) was divided by the DAPI signal profile (Fig. 2F) for adjustment. The adjusted Sox9 profile (Fig. 2G) was then used for the following “change point analysis”.

Excessive chondrogenesis in the blastema was assessed by the adjusted Sox9 profile from distal to proximal (Fig. S1). If no significant chondrogenesis occurred in the blastema, no change point would be detected (Fig. S1A). In contrast, if excessive chondrogenesis occurred (i.e., it invaded the blastema), a change point would be detected at a certain level along the PD axis. The more significant the excessive chondrogenesis, the more proximal the change point would be detected (Fig. S1B, C). Using the procedure described above, we compared the extent of chondrogenesis in the blastemas of newts, tadpoles, and froglets based on these same criteria.

### A change point was specifically detected in the froglet blastema: excessive chondrogenesis seems to initiate as early as the medium bud (MB) stage

Double immunofluorescence staining for Sox9 and Msx1/2 was used to visualize the spatial localization of differentiating early chondrocyte and blastema cells, respectively, in the sections of newt, tadpole, and froglet blastemas (Fig. 3). In the MB blastemas of a newt or tadpole, Sox9^+^ cells were rarely found in the rectangle region of interest (ROI) (Fig. 3A’, B’). In contrast, Sox9^+^ cells were distributed in the proximal part of the ROI in the MB blastema of the froglet (Fig. 3C’). At the early bud (EB) stage, none of newt (Fig. S2A’), tadpole (Fig. S2C’) or froglet blastemas (Fig. S3A’) appeared to have sufficient Sox9 signals indicative of chondrocytes in the rectangular ROI. Change point analysis produced consistent results with a change point detected only in the froglet blastema at the MB stage (Fig. 3C-C’’; the change point value was 0.82), among the representative blastemas of newts, tadpoles, and froglets at EB and MB stages shown in Fig. 3, Fig. S2 and Fig. S3.

In all examined blastema samples, no change point was detected in newt and tadpole blastemas at the EB and MB stages (Fig. 4A, top and middle). In the froglet blastemas, a change point was detected in 33% of the blastemas at the MB stage, while no change point was detected in EB blastemas (Fig. 4A, bottom). These results suggest that the first sign of excessive chondrogenesis in froglet blastema emerges at the MB stage, and that the fate of froglet blastema, with respect to cartilage differentiation, diverges from that of the newt and tadpole blastema at the MB stage at the latest.

**Fig. 4.**
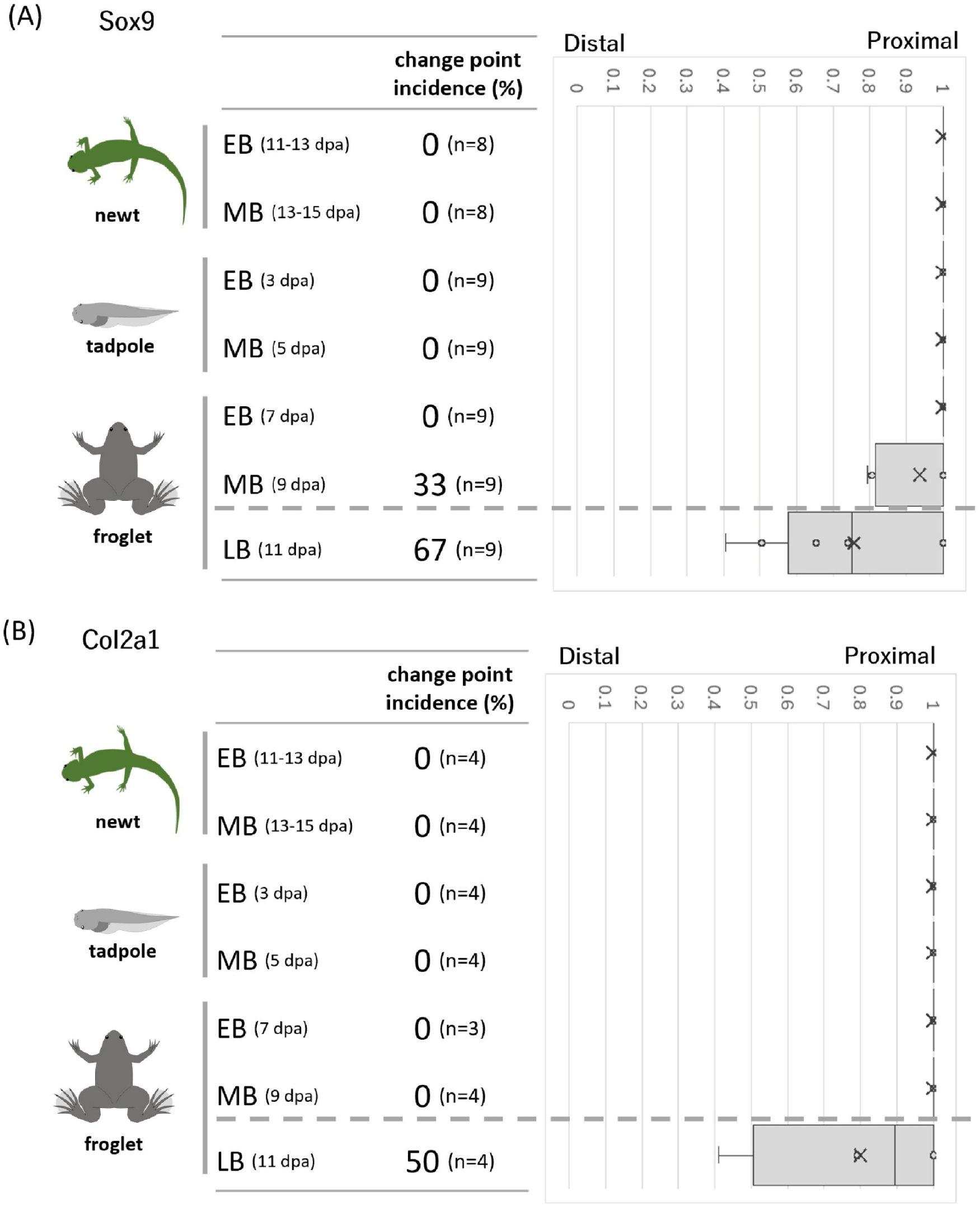
Incidence of change point detection and its value in newt, tadpole and froglet blastemas using anti-Sox9 or anti-Col2a1 antibody. Incidence (%) of change point detection and box-and-whisker plot showing change point value using anti-Sox9 (A) or anti-Col2a1 antibody (B). In each sample group, vertical lines indicate the minimum, first quartile, median, third quartile, and maximum values, from left to right. X, the mean value; n, number of blastemas used for change point analysis.

To examine whether excessive chondrogenesis in froglet blastemas progresses into a more significant state as the blasetmas grow, we conducted a change point analysis on the late bud (LB) blastemas, as well. In the froglet LB blastemas, a change point was detected at a higher incidence (67%) than in the MB blastemas, and the change point values, both as mean and median, shifted more distally (Fig. 4A, bottom). Concomitantly, many Sox9^+^ cells were broadly distributed in the rectangular ROI, except for the distal-most area, in the froglet blastema at the LB stage, as shown in Fig. S3C’. These results suggest that chondrogenesis in froglet blastemas progresses to a more excessive state at the LB stage.

Sox9 activates the transcription of genes encoding collagen type II (Col2a1), a typical cartilage marker, as one of its direct target genes (Egawa et al., 2014; Kozhemyakina et al., 2015). If Sox9^+^ early chondrocytes differentiate into Col2a1^+^ cartilage in the froglet blastema, the spatial pattern of Sox9 should be recapitulated by that of Col2a1 at later stage, resulting in the detection of a change point using an anti-Col2a1 antibody. To test this hypothesis, we conducted immunofluorescence and change point analysis using an anti-Col2a1 antibody instead of an anti-Sox9 antibody. Note that we performed immunofluorescence for Msx1/2 on the adjacent sections, rather than the same sections as for Col2a1 (Fig. S4 and S5), since immunostaining for Col2a1 requires proteinase K treatment for antigen retrieval (see Materials and Methods for details). Col2a1^+^ cells were found in the proximal-most area of the rectangular ROI in the froglet blastema at the LB stage (Fig. S5C’), similarly to the Sox9^+^ chondrocytes found in the ROI of the froglet blastema at the MB stage (Fig. 3C’). In newt, tadpole, and froglet blastemas at the EB and MB stages, however, such Col2a1^+^ cells were not found in the rectangular ROI (Fig. S4A’-D’ and Fig. S5A’, B’). Consistent with these results, a change point was detected only in the froglet blastemas at the LB stage (Fig. 4B, bottom). This result suggests that the sign of excessive chondrogenesis indicated by the spatial distribution of Sox9 actually induces excessive chondrogenesis (cartilage differentiation), which is underpinned by the distribution of Col2a1 in the subsequent stage of blastemas.

### How limb regeneration in a froglet can be improved by repressing excessive chondrogenesis

Ohgo et al. (2010) raised the possibility that premature and excessive chondrogenesis may prevent pattern formation as well as muscle formation in the limb blastema of froglets, based on expression analysis on *hoxa11*, *hoxa13*, and *sox9*. Aztekin et al. (2021) reported the negative impact of developing cartilage on limb regeneration from a different perspective, using regeneration-restricted limbs in *X. laevis* tadpoles. Noggin (Nog), secreted by developing cartilage, blocks the formation of functional apical epidermis in the blastema, and FGF10 overrides this negative effect by suppressing chondrogenesis (Aztekin et al., 2021). While these studies suggest that appropriate inhibition of excessive chondrogenesis in the froglet blastema could improve limb regeneration, it was unclear when and how excessive chondrogenesis initiates in the froglet blastema. Our work revealed that the first sign of excessive chondrogenesis, based on Sox9 distribution in froglet, is observed by the MB stage at the latest. This finding provides a specific target time point: we should suppress excessive chondrogenesis, likely around the MB stage, and interrupt the subsequent progression of excessive chondrogenesis to improve limb regeneration capacity.

To suppress chondrogenesis at the appropriate time point, what kind of method should we adopt? Conventional knockout of the *sox9* gene using genome editing would not be a good choice, as *sox9* plays an essential role in multiple aspects of embryonic development. In fact, CRISPR-based *sox9* knockout *X. tropicalis* individuals (crispants) exhibit various defects at tadpole stages due to defects in neural crest and other tissue progenitor formation (Hossain et al., 2023). To improve limb regeneration in froglets, we expect that chondrogenesis should be suppressed to an adequate level, but should not be completely eliminated. One potential approach is the overexpression of a dominant-negative form of the *sox9* gene. It is feasible to design a dominant-negative form of *sox9* based on loss-of-function studies on other Sox family genes (e.g., Kishi et al., 2000; Mizuseki et al., 1998). Temporally and spatially restricted induction of gene expression is now possible in *Xenopus* froglets using a heat-shock-inducible system (Kawasumi-Kita et al., 2015; Matsubara et al., 2023b). The Tet-on system also appears to be promising for this purpose in *X. laevis* (Sterner et al., 2019). This system requires a blastema-specific promoter/enhancer, and *prrx1* limb-specific enhancer would fulfill this requirement with its specific activation in the limb blastema (Suzuki et al., 2007). When the dominant-negative form of *sox9* is overexpressed at a specific time point, we need to assess the attenuation of Sox9 function in the froglet blastema. Since our study suggests that immunofluorescence for Col2a1 and subsequent change point analysis follows the corresponding data for Sox9 (Fig. S5 and Fig, 4), Col2a1 would be a good marker to check and optimize Sox9 function attenuation. It is very intriguing to examine whether attenuation of Sox9 function recover pattern formation and muscle formation during limb regeneration in froglets in the near future.

It is still largely unknown why excessive chondrogenesis specifically initiates in froglet blastemas. Single-cell RNA-seq analysis, along with transplant experiments, revealed that cartilage differentiation programs in froglet limb regeneration are molecularly distinct from those in tadpole limb development (Lin et al., 2021). This finding implies that identifying the initiation factor(s) that trigger froglet-specific cartilage differentiation is the next important task. While FGF10 suppresses chondrogenesis in regenerating limb (Aztekin et al., 2021), it is unlikely that the simple lack of *fgf10* expression causes excessive chondrogenesis in froglet blastemas. *fgf10*, *fgf8*, and *msx1* are expressed in froglet blastemas (Suzuki et al., 2005). Additionally, the fact that *dusp6* expression, which reflects the activity of ETS transcription factors induced by FGF signaling, is also observed in froglet blastemas (Tada et al., 2023), suggests the presence of an FGF10/FGF8 feedback loop. However, another important feedback loop in limb development, the FGF/Shh loop (Verheyden and Sun, 2008) appears to be absent in froglet blastemas, as *shh* is not expressed in froglet blastemas (Endo et al., 2000; Yakushiji et al., 2007). The absence of the FGF/Shh loop may indirectly influence excessive chondrogenesis in froglet blastemas by modifying FGF activity, which normally suppresses chondrogenesis. The spatio-temporal signs of excessive chondrogenesis provided by this study, especially the distribution of Sox9 in froglet blastemas, could offer a clue to identifying the initiation factor(s) responsible for triggering froglet-specific cartilage differentiation. The initiation factor(s) should coincide with or precede the spatio-temporal signs of excessive chondrogenesis. We expect that elucidating the froglet-specific transcriptional regulation of *sox9*, with reference to anti-chondrogenic factors such as FGF activity in the blastema, will be especially important in unraveling the mechanisms of excessive chondrogenesis in froglet blastemas. This may lead to effective suppression of chondrogenesis and the recovery of patterned limb regeneration in froglets.

### Conclusions

In this study, we established a simplified and reproducible procedure to evaluate excessive chondrogenesis in regenerating limb blastema using immunofluorescence for cartilage markers (Sox9 or Col2a1) and quantitative image analysis with ImageJ. This procedure, which we refer to as change point analysis, enables to interspecific (urodele versus *Xenopus*) comparisons as well as intraspecific (tadpole versus froglet in *Xenopus*) comparisons to examine whether excessive chondrogenesis specifically occurs in hypomorphic regeneration in froglets. As a result, change point analysis for Sox9 detected signs of excessive chondrogenesis not in newt or tadpole blastemas but in froglet blastemas as early as at the medium bud (MB) stage. Furthermore, the sign of excessive chondrogenesis (change point) progressed more distally as the froglet blastemas grew to the late bud (LB) stage. These findings delineated the spatio-temporal progression pattern of excessive chondrogenesis in froglet blastemas and provided an important clue to elucidating the initiation mechanisms of froglet-specific excessive chondrogenesis. Since premature and excessive chondrogenesis has been thought to exert negative effects on pattern formation in limb regeneration of froglets (Ohgo et al., 2010), adequate inhibition of the initiation of excessive chondrogenesis in froget blastemas may lead to the recovery of patterned limb regeneration.

## ACKNOWLEDGMENTS

This work was supported by the NIBB Collaborative Research Program to HY (19-451, 20-436, 21-307, 21-401, 22NIBB428, 22NIBB501, 23NIBB406, 23NIBB501, 24NIBB453, 24NIBB523) and the Joint Research of the Exploratory Research Center on Life and Living Systems (ExCELLS) (ExCELLS program No, 24EXC355) to HY. This work was supported by JSPS KAKENHI Grant Number JP22H04926, Grant-in-Aid for Transformative Research Areas ― Platforms for Advanced Technologies and Research Resources “Advanced Bioimaging Support”. This work was supported by the National Bio-Resource Project (NBRP) of the MEXT, Japan. Iberian ribbed newts (*Pleurodeles waltl*) were provided by Hiroshima University Amphibian Research Center (RRID: SCR_019015) through National BioResource Project (NBRP) of MEXT. We acknowledge Shared Facility Center for Science and Technology, Hirosaki University (SFCST) for confocal microscope observation by FV3000 (Olympus). We thank Dr. Ikuo Uchiyama for support in gene expression analysis. We thank Dr. Michiko Sasabe, Dr. Shigenori Nonaka and Misako Saida for their help in using confocal microscopy. We thank Dr. Makoto Suzuki for advices on immunostaining of the blastema with ant-Msx1/2 antibody. We thank Drs. Haruka Matsubara and Tetsuya Endo for advices on immunostaining of the blastema with ant-Col2a1 antibody. We thank Drs. Haruka Matsubara and Kiyokazu Agata for providing *Xenopus laevis col2a1* cDNA. We thank Dr. Atsuo Nishino and members of Nishino laboratory for their support for histological staining. We thank Dr. Toshinori Hayashi, Dr. Haruka Matsubara and Arisa Anami for their multiple helps in start using Iberian ribbed newts in Yokoyama laboratory.

## REFERENCES

Aztekin, C., Hiscock, T.W., Gurdon, J., Jullien, J., Marioni, J., Simons, B.D., 2021. Secreted inhibitors drive the loss of regeneration competence in *Xenopus* limbs. Development 148. dev199158. 10.1242/dev.199158.

Dent, J.N., 1962. Limb regeneration in larvae and metamorphosing individuals of the South African clawed toad. J Morphol 110, 61–77.

Egawa, S., Miura, S., Yokoyama, H., Endo, T., Tamura, K., 2014. Growth and differentiation of a long bone in limb development, repair and regeneration. Dev Growth Differ 56, 410–424.

Endo, T., Tamura, K., Ide, H., 2000. Analysis of gene expressions during *Xenopus* forelimb regeneration. Dev Biol 220, 296–306.

Hayashi, T., Yokotani, N., Tane, S., Matsumoto, A., Myouga, A., Okamoto, M., Takeuchi, T., 2013. Molecular genetic system for regenerative studies using newts. Dev Growth Differ 55, 229–236.

Hossain, N., Igawa, T., Suzuki, M., Tazawa, I., Nakao, Y., Hayashi, T., Suzuki, N., Ogino, H., 2023. Phenotype-genotype relationships in *Xenopus sox9* crispants provide insights into campomelic dysplasia and vertebrate jaw evolution. Dev Growth Differ 65, 481–497.

Iten, L.E., Bryant, S.V., 1973. Forelimb regeneration from different levels of amputation in the newt,Notophthalmus viridescens: Length, rate, and stages. Wilhelm Roux Arch Entwickl Mech Org 173, 263–282.

Joven, A., Elewa, A., Simon, A., 2019. Model systems for regeneration: salamanders. Development 146. dev167700. 10.1242/dev.167700.

Kawasumi-Kita, A., Hayashi, T., Kobayashi, T., Nagayama, C., Hayashi, S., Kamei, Y., Morishita, Y., Takeuchi, T., Tamura, K., Yokoyama, H., 2015. Application of local gene induction by infrared laser-mediated microscope and temperature stimulator to amphibian regeneration study. Dev Growth Differ 57, 601–613.

Killick, R., Eckley, I.A., 2014. changepoint: An R Package for Changepoint Analysis. Journal of Statistical Software 58, 1–19.

Kishi, M., Mizuseki, K., Sasai, N., Yamazaki, H., Shiota, K., Nakanishi, S., Sasai, Y., 2000. Requirement of Sox2-mediated signaling for differentiation of early Xenopus neuroectoderm. Development 127, 791–800.

Kozhemyakina, E., Lassar, A.B., Zelzer, E., 2015. A pathway to bone: signaling molecules and transcription factors involved in chondrocyte development and maturation. Development 142, 817–831.

Lin, T.Y., Gerber, T., Taniguchi-Sugiura, Y., Murawala, P., Hermann, S., Grosser, L., Shibata, E., Treutlein, B., Tanaka, E.M., 2021. Fibroblast dedifferentiation as a determinant of successful regeneration. Dev Cell 56, 1541–1551.e1546.

Matsubara, H., Inoue, T., Agata, K., 2023a. ECM degradation in the stump region induced by Fgf during functional joint regeneration in frogs. bioRxiv. 10.1101/2023.08.04.551975.

Matsubara, H., Kawasumi-Kita, A., Nara, S., Yokoyama, H., Hayashi, T., Takeuchi, T., 2023b. Appendage-restricted gene induction using a heated agarose gel for studying regeneration in metamorphosed *Xenopus laevis* and *Pleurodeles waltl*. Dev Growth Differ 65, 86–93.

Matsuda, H., Yokoyama, H., Endo, T., Tamura, K., Ide, H., 2001. An epidermal signal regulates *Lmx-1* expression and dorsal-ventral pattern during *Xenopus* limb regeneration. Dev Biol 229, 351–362.

Matsunami, M., Suzuki, M., Haramoto, Y., Fukui, A., Inoue, T., Yamaguchi, K., Uchiyama, I., Mori, K., Tashiro, K., Ito, Y., Takeuchi, T., Suzuki, K.T., Agata, K., Shigenobu, S., Hayashi, T., 2019. A comprehensive reference transcriptome resource for the Iberian ribbed newt *Pleurodeles waltl*, an emerging model for developmental and regeneration biology. DNA Res 26, 217–229.

Mizuseki, K., Kishi, M., Shiota, K., Nakanishi, S., Sasai, Y., 1998. SoxD: an essential mediator of induction of anterior neural tissues in Xenopus embryos. Neuron 21, 77–85.

Nieuwkoop, P.D., Faber, J., 1994. Normal Table of Xenopus laevis (Daudin). Garland Publishing, New York.

Ohgo, S., Itoh, A., Suzuki, M., Satoh, A., Yokoyama, H., Tamura, K., 2010. Analysis of *hoxa11* and *hoxa13* expression during patternless limb regeneration in *Xenopus*. Dev Biol 338, 148–157.

Otsuki, L., Tanaka, E.M., 2022. Positional Memory in Vertebrate Regeneration: A Century’s Insights from the Salamander Limb. Cold Spring Harb Perspect Biol 14. 10.1101/cshperspect.a040899.

Robinson, H., Allenby, K., 1974. The effect of nerve growth factor on hindlimb regeneration in Xenopus laevis froglets. J Exp Zool 189, 215–226.

Satoh, A., Ide, H., Tamura, K., 2005. Muscle formation in regenerating *Xenopus* froglet limb. Dev Dyn 233, 337–346.

Sterner, Z.R., Rankin, S.A., Wlizla, M., Choi, J.A., Luedeke, D.M., Zorn, A.M., Buchholz, D.R., 2019. Novel vectors for functional interrogation of *Xenopus* ORFeome coding sequences. Genesis 57, e23329.

Stocum, D.L., 1995. Regeneration of amphibian limbs, Wound Repair, Regeneration and Artificial Tissues. R. G. Landes, pp. 51–79.

Stoick-Cooper, C.L., Moon, R.T., Weidinger, G., 2007. Advances in signaling in vertebrate regeneration as a prelude to regenerative medicine. Genes Dev 21, 1292–1315.

Straube, W.L., Tanaka, E.M., 2006. Reversibility of the differentiated state: regeneration in amphibians. Artif Organs 30, 743–755.

Suzuki, M., Satoh, A., Ide, H., Tamura, K., 2005. Nerve-dependent and -independent events in blastema formation during *Xenopus* froglet limb regeneration. Dev Biol 286, 361–375.

Suzuki, M., Satoh, A., Ide, H., Tamura, K., 2007. Transgenic *Xenopus* with *prx1* limb enhancer reveals crucial contribution of MEK/ERK and PI3K/AKT pathways in blastema formation during limb regeneration. Dev Biol 304, 675–686.

Suzuki, M., Yakushiji, N., Nakada, Y., Satoh, A., Ide, H., Tamura, K., 2006. Limb regeneration in *Xenopus laevis* froglet. ScientificWorldJournal 6 Suppl 1, 26–37.

Tada, R., Higashidate, T., Amano, T., Ishikawa, S., Yokoyama, C., Kobari, S., Nara, S., Ishida, K., Kawaguchi, A., Ochi, H., Ogino, H., Yakushiji-Kaminatsui, N., Sakamoto, J., Kamei, Y., Tamura, K., Yokoyama, H., 2023. The *shh* limb enhancer is activated in patterned limb regeneration but not in hypomorphic limb regeneration in *Xenopus laevis*. Dev Biol 500, 22–30.

Tschumi, P.A., 1957. The growth of the hindlimb bud of *Xenopus laevis* and its dependence upon the epidermis. J Anat 91, 149–173.

Verheyden, J.M., Sun, X., 2008. An Fgf/Gremlin inhibitory feedback loop triggers termination of limb bud outgrowth. Nature 454, 638–641.

Yakushiji, N., Suzuki, M., Satoh, A., Sagai, T., Shiroishi, T., Kobayashi, H., Sasaki, H., Ide, H., Tamura, K., 2007. Correlation between *Shh* expression and DNA methylation status of the limb-specific *Shh* enhancer region during limb regeneration in amphibians. Dev Biol 312, 171–182.

Yoshida, K., Urase, K., Takahashi, J., Ishii, Y., Yasugi, S., 1996. Mucus-associated antigen in epithelial cells of the chicken digestive tract: developmental change in expression and implications for morphogenesis-function relationships. Dev. Growth Differ. 38, 185–192.

