## Supplemental Figures for "Excessive chondrogenesis in the blastema initiates during hypomorphic limb regeneration in *Xenopus* froglet, but not during patterned limb regeneration in *Xenopus* tadpoles and newts"

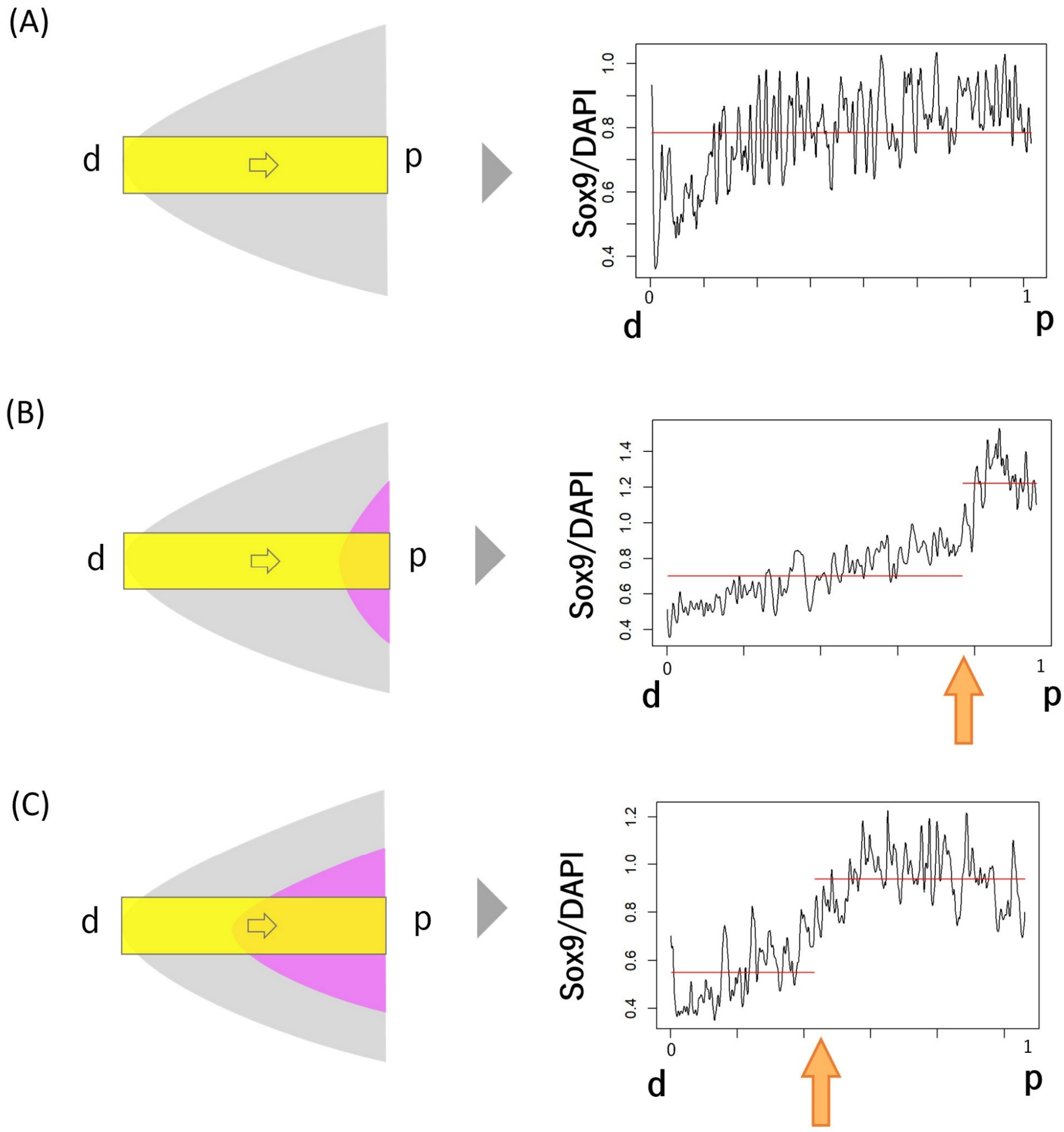


**Fig. S1. Expected results of change point analysis in blastemas with different extents of excessive chondrogenesis.** (A) No significant chondrogenesis in a blastema (left). No change point will be detected in Sox9/DAPI plot profile (right). (B) Excessive chondrogenesis in a blastema (left). Change point will be detected in the profile (right). (C) More severe excessive chondrogenesis in a blastema (left). Change point will be detected at more distal site (at smaller value) in the profile (right). An orange arrow indicates a detected change point. d, distal; p, proximal.

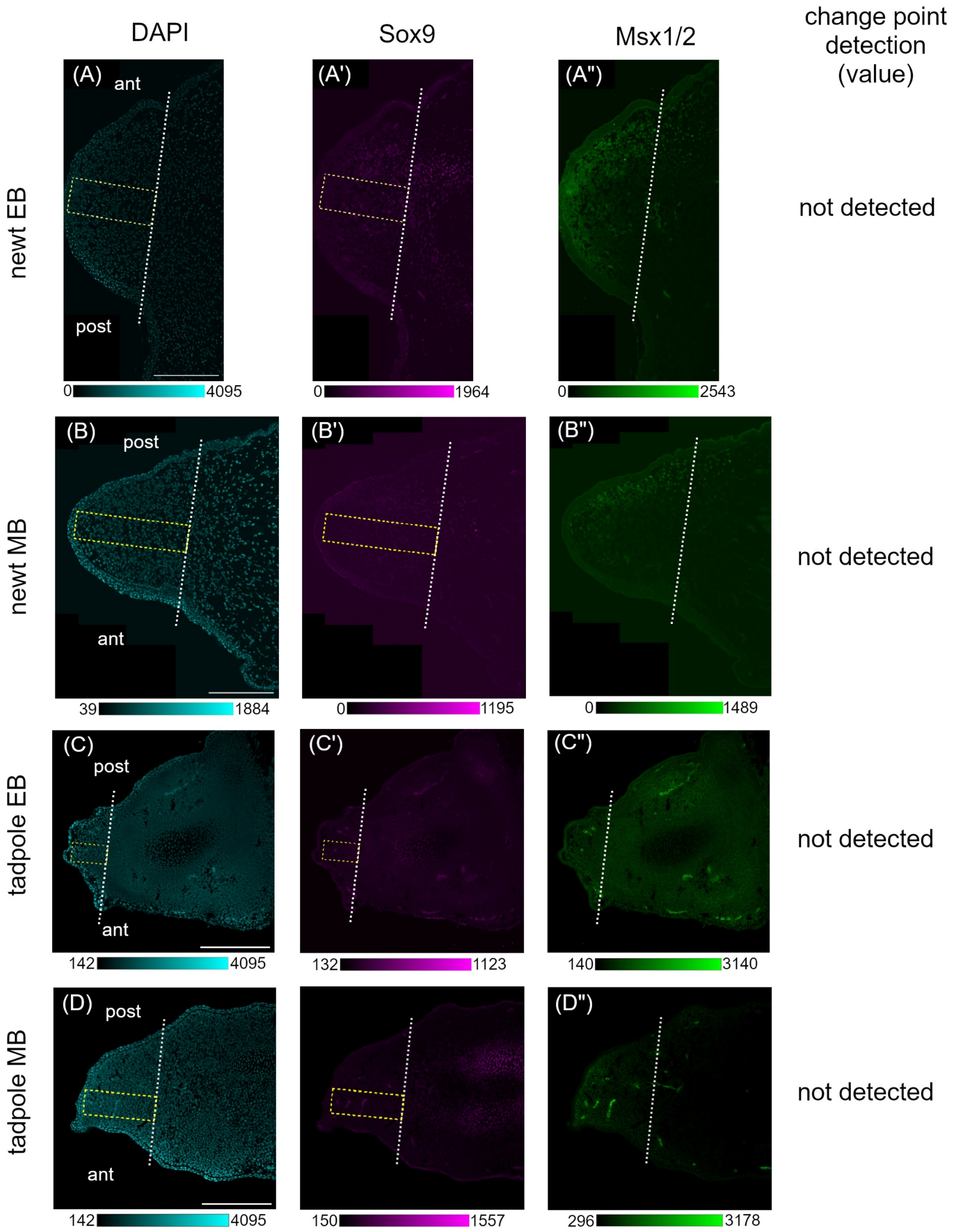


**Fig. S2. Representative immunofluorescence images of EB and MB blastemas where change point (Sox9) was not detected in newts and tadpoles.** Fluorecence images of a longitudinal section of newt EB (A-A''), newt MB (B-B'', identical to Fig. 3A-A''), tadpole EB (C-C'') and tadpole MB blastemas (D-D'', identical to Fig. 3B-B''). A yellow rectangle indicates the region of interest (ROI) for analysis. A dotted white line indicates the amputation level. A calibration bar, created using ImageJ, represents the range of grey values in each image. ant, anterior; post, posterior. Bars = 400 µm for (A, B) and 200 µm for (C, D).

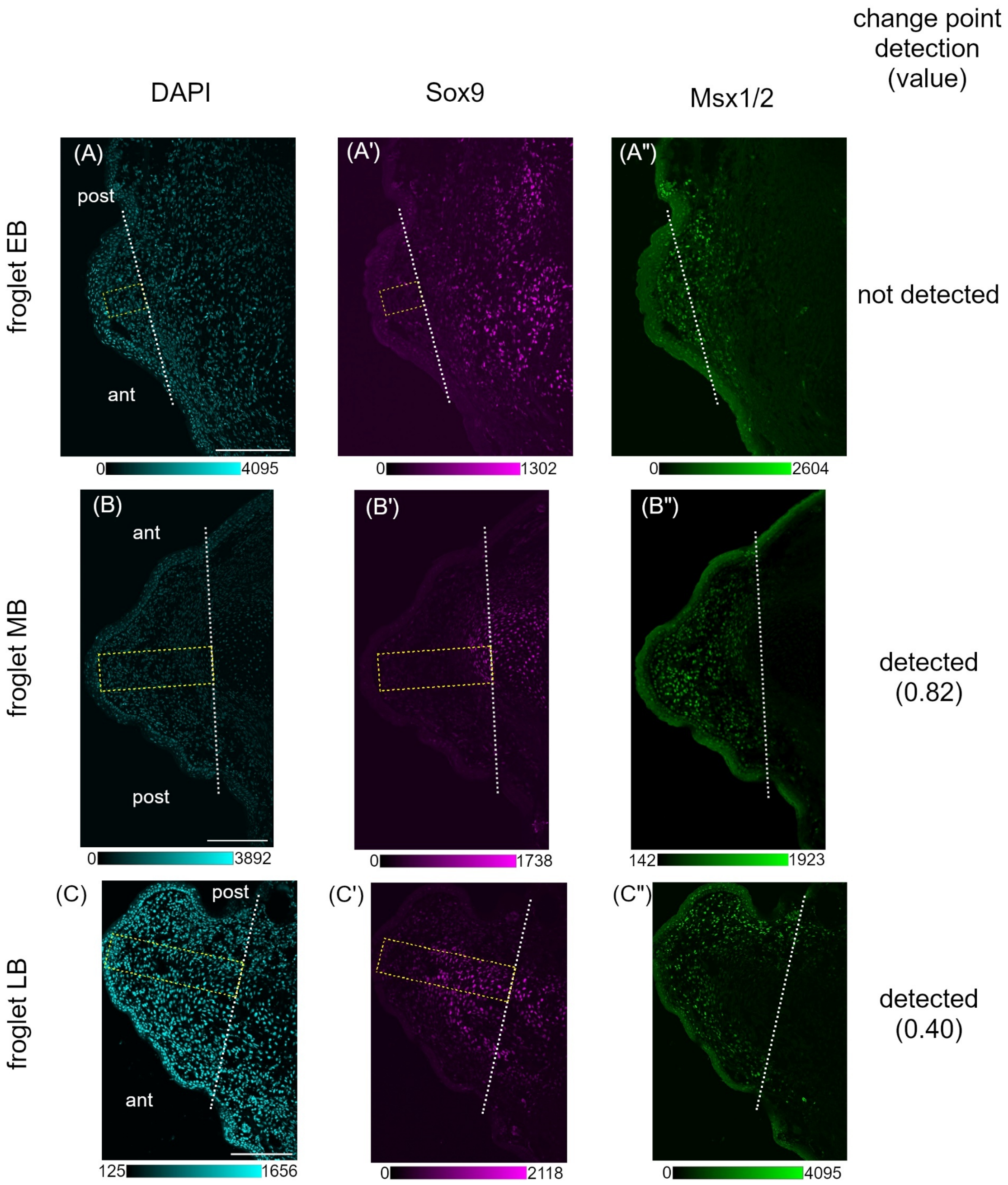


**Fig. S3. Representative immunofluorescence images of froglet blastemas where change point (Sox9) was detected at MB and LB stages.** Fluorecence images of a longitudinal section of EB (A-A''), MB (B-B'', identical to Fig. 3C-C''), LB blastemas (C-C'', identical to Fig. 3D-D''). A yellow rectangle indicates the region of interest (ROI) for analysis. A dotted white line indicates the amputation level. A calibration bar, created using ImageJ, represents the range of grey values in each image. ant, anterior; post, posterior. Bars = 200 μm.

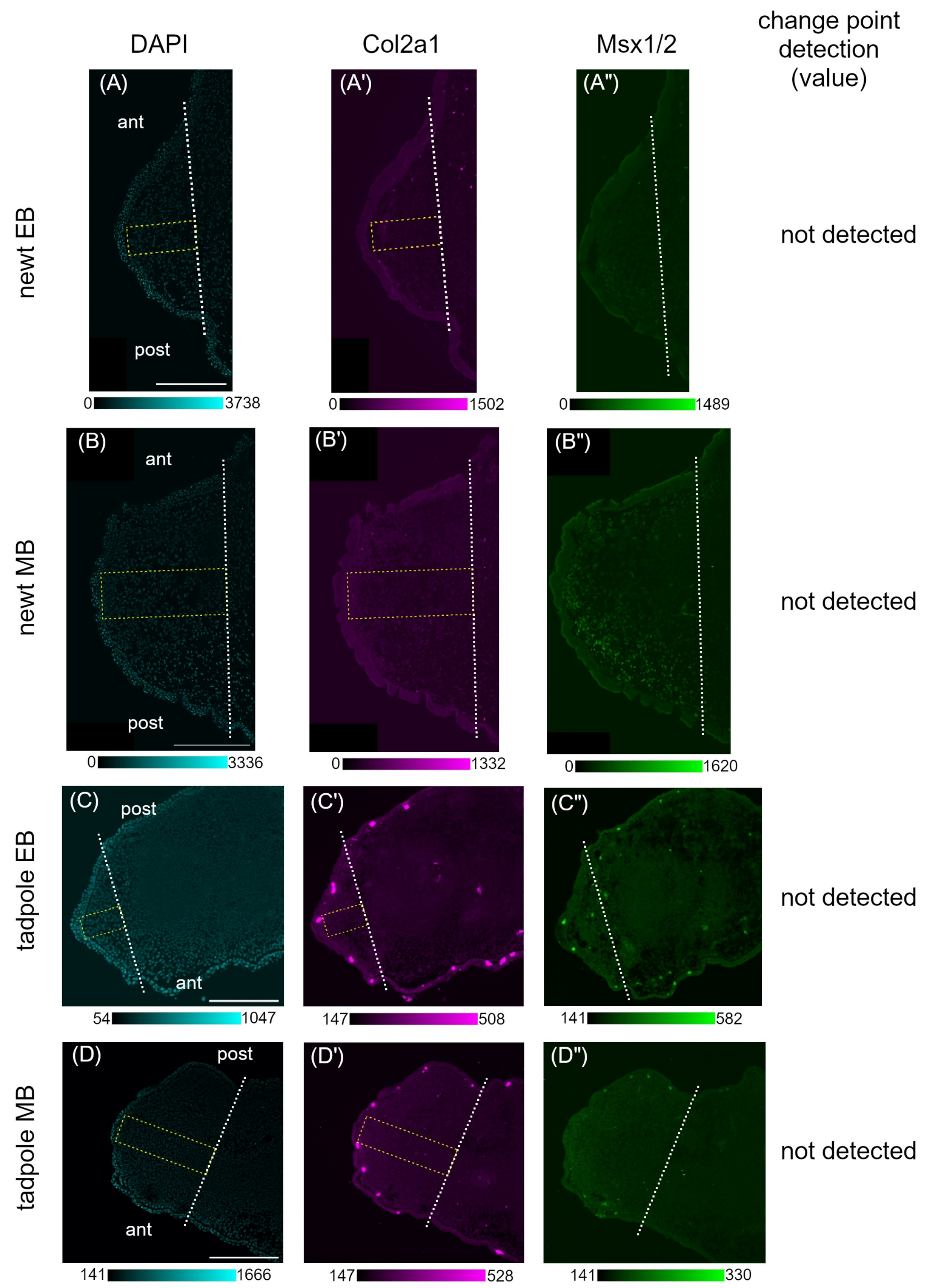


**Fig. S4. Representative immunofluorescence images of EB and MB blastemas where change point (Col2a1) was not detected in newts and tadpoles.** Fluorecence images of adjacent longitudinal sections from newt EB (A-A''), newt MB (B-B''), tadpole EB (C-

C'') and tadpole MB blastemas (D-D''). Note that the Msx1/2 images (A'', B'', C'', D'') were captured from adjacent sections, not the same one, as the DAPI (A, B, C, D) and Col2a1 images (A', B', C', D'), respectively. A yellow rectangle indicates the region of interest (ROI) for analysis. A dotted white line indicates the amputation level. A calibration bar, created using ImageJ, represents the range of grey values in each image. ant, anterior; post, posterior. Bars = 400 μm for (A, B) and 200 μm for (C, D).

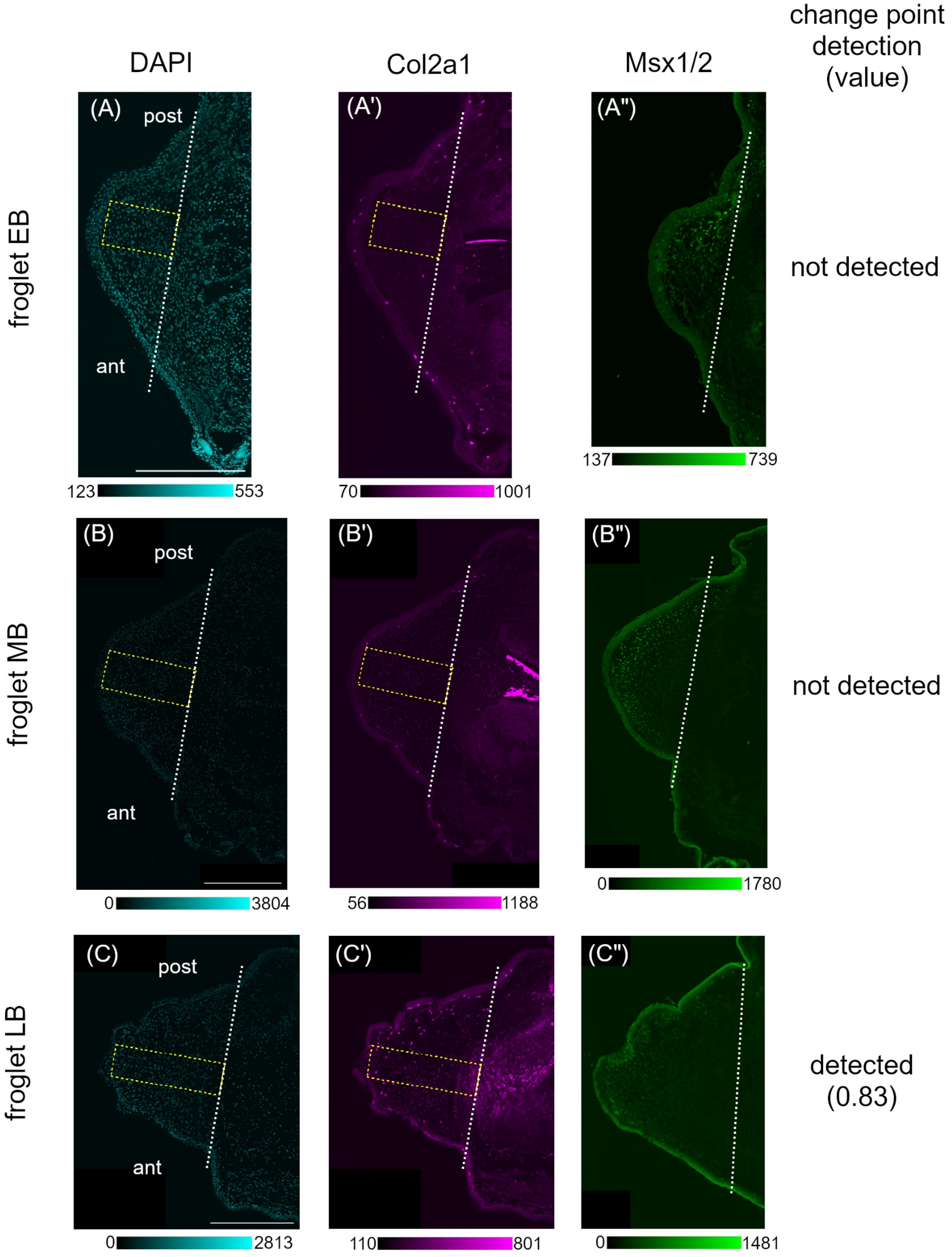


**Fig. S5. Representative immunofluorescence images of froglet blastemas where change point (Col2a1) was detected at LB stage.** Fluorecence images of adjacent longitudinal sections from EB (A-A''), MB (B-B'') and LB blastemas (C-C''). Note that the Msx1/2 images (A'', B'', C'') were captured from adjacent sections, not the same one, as the DAPI (A, B, C) and Col2a1 images (A', B', C'), respectively. A yellow rectangle indicates the region of interest (ROI) for analysis. A dotted white line indicates the amputation level. A calibration bar, created using ImageJ, represents the range of grey values in each image. ant, anterior; post, posterior. Bars = 400 μm.
